# A competence-regulated toxin-antitoxin system in *Haemophilus influenzae*

**DOI:** 10.1101/330480

**Authors:** Hailey Findlay Black, Scott Mastromatteo, Sunita Sinha, Rachel L. Ehrlich, Corey Nislow, Joshua Chang Mell, Rosemary J. Redfield

## Abstract

Natural competence allows bacteria to respond to environmental and nutritional cues by taking up free DNA from their surroundings, thus gaining nutrients and genetic information. In the Gram-negative bacterium *Haemophilus influenae*, the DNA uptake machinery is induced by the CRP and *Sxy* transcription factors in response to lack of preferred carbon sources and nucleotide precursors. Here we show that *HI0659*—which is absolutely required for DNA uptake— encodes the antitoxin of a competence-regulated toxin-antitoxin operon (‘*toxTA’*), likely acquired by horizontal gene transfer from a *Streptococcus* species. Deletion of the toxin restores uptake to the antitoxin mutant. In addition to the expected Sxy-and CRP-dependent-competence promoter, transcript analysis using RNA-seq identified an internal antitoxin-repressed promoter whose transcription starts within *toxT* and will yield nonfunctional protein. We present evidence that the most likely effect of unopposed toxin expression is non-specific cleavage of mRNAs and arrest or death of competent cells in the culture, and we show that the toxin gene has been inactivated by deletion in many *H. influenzae* strains. We suggest that this competence-regulated toxin-antitoxin system may facilitate downregulation of protein synthesis and recycling of nucleotides under starvation conditions, or alternatively be a simple genetic parasite.

## INTRODUCTION

Toxin-antitoxin systems are bacterial gene pairs that were originally discovered on plasmids, where they function to promote plasmid persistence by killing any daughter cells that have not inherited the plasmid. Typically, one gene of the pair encodes a relatively stable toxic protein that blocks cell growth, and the other encodes a labile antitoxin (RNA or protein) that blocks the toxin’s activity and regulates its expression (Yamaguchi *et al.,* 2011, Goeders and Van Melderen, 2014). Similar toxin-antitoxin gene pairs have been discovered on many bacterial chromosomes, where they are thought to be relatively recent introductions that in some cases have been co-opted to regulate cellular functions (Van Melderen and Saavedra de Bast, 2009). Here we describe one such system, which is induced in naturally competent cells and whose unopposed toxin completely prevents DNA uptake and transformation.

Many bacteria are naturally competent, able to take up DNA from their surroundings and—when sequence similarity allows—recombine it into their genomes (Ambur *et al.*, 2016, Johnston *et al.*, 2014, Mell and Redfield, 2014). In most species, this DNA uptake is tightly controlled, with protein machinery specified by a set of co-regulated chromosomal genes induced in response to diverse cellular signals. In addition to components of the DNA-uptake machinery, competence-regulon genes encode proteins that translocate DNA across the inner membrane, proteins that facilitate recombination, and proteins of unknown function. *Haemophilus influenzae* has an unusually small and well-defined competence regulon (26 genes in 13 operons) induced by signals of energy and nucleotide scarcity (Redfield *et al*., 2005, Sinha *et al*., 2012). Induction of these genes begins in response to depletion of phosphotransferase sugars, when rising levels of cyclic AMP (cAMP) first stimulate transcription of genes regulated by the transcriptional activator CRP. One of these induced genes encodes the competence-specific transcriptional activator Sxy, but efficient translation of *sxy* mRNA occurs only when purine pools are also sufficiently depleted (Macfadyen *et al*., 2001; Sinha *et al*., 2013). Sxy then acts with CRP at the promoters of competence genes, stimulating their expression and leading to DNA uptake and natural transformation. These competence promoters are distinguished by the presence of ‘CRP-S’ sites (formerly called CRE sites), variants of standard CRP sites that depend on both CRP and Sxy for activation (Cameron and Redfield, 2006).

All but one of the fifteen *H. influenzae* genes needed for DNA uptake encode typical competence proteins– membrane-associated proteins homologous to known components of the Type IV pilus-based DNA uptake machinery present in nearly all known naturally competent species (Johnston *et al*., 2014). The one exception is *HI0659*, which instead encodes a 98 amino acid cytoplasmic protein with no similarity to known DNA uptake proteins. It shares a competence-inducible CRP-S promoter with an upstream gene encoding another short cytoplasmic protein (*HI0660*, 119 aa) (**Fig. 1, top**). Although a knockout of *HI0659* eliminates detectable DNA uptake and transformation, a knockout of *HI0660* has no effect (Sinha *et al.* 2012).

**Figure 1:**
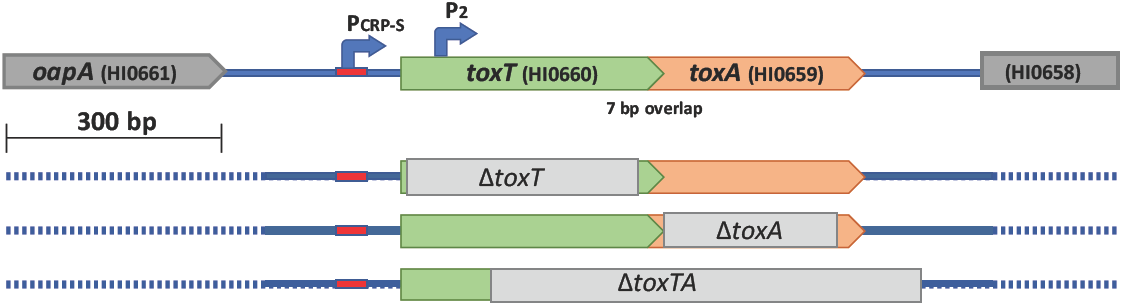
Structure of wildtype and mutant *toxTA* genes. Top line: structure of the wildtype *toxTA* operon in strain KW20. Lower lines: light grey bars indicate segments deleted in Δ*toxT*, Δ*toxA*, and Δ*toxT*A mutants.

Here we show that *HI0660* and *HI0659* comprise a horizontally transferred operon that encodes a toxin-antitoxin pair, and that expression of the toxin in the absence of the antitoxin completely prevents DNA uptake and transformation. Surprisingly, this unopposed toxin expression has only slight effects on induction of competence genes, and on cell growth and viability. The HI0660 toxin is unusual in that its overexpression in the absence of HI0659 is not lethal to cells growing in rich medium, which may be explained by our observation that transcription in the absence of HI0659 occurs mainly from a second internal promoter that would not produce functional protein.

## RESULTS

### HI0659 and HI0660 act as a toxin-antitoxin system

Our original analyses of competence-induced genes did not identify any close homologs of *HI0659* or *HI0660* (Redfield *et al*. 2005, Sinha *et al*. 2012). However recent database searches and examination of BLAST results revealed that these genes’ products resemble proteins in the Type II toxin/antitoxin families, which typically occur in similar two-gene operons. If *HI0660* and *HI0659* do encode a toxin-antitoxin pair, then *ΔHI0659*’s DNA uptake defect would likely be caused by unopposed expression of a *HI0660*-encoded toxin protein that prevents DNA uptake, so knocking out this toxin gene should restore competence to the *HI0659* (antitoxin^-^) mutant. We tested this by constructing an *HI0660/HI0659* double mutant (Fig. 1) and examining its ability to be transformed with antibiotic-resistant chromosomal DNA. The double mutant had normal transformation (**Fig. 2**), showing that mutation of *HI0660* suppresses the competence defect of an *HI0659* mutant, and also that neither *HI0660* nor *HI0659* is directly needed for the development of competence. This supported the postulated antitoxin function of *HI0660*, so we named the *HI0660* and *HI0659* genes *toxT* (toxin) and *toxA* (antitoxin) respectively.

**Figure 2:**
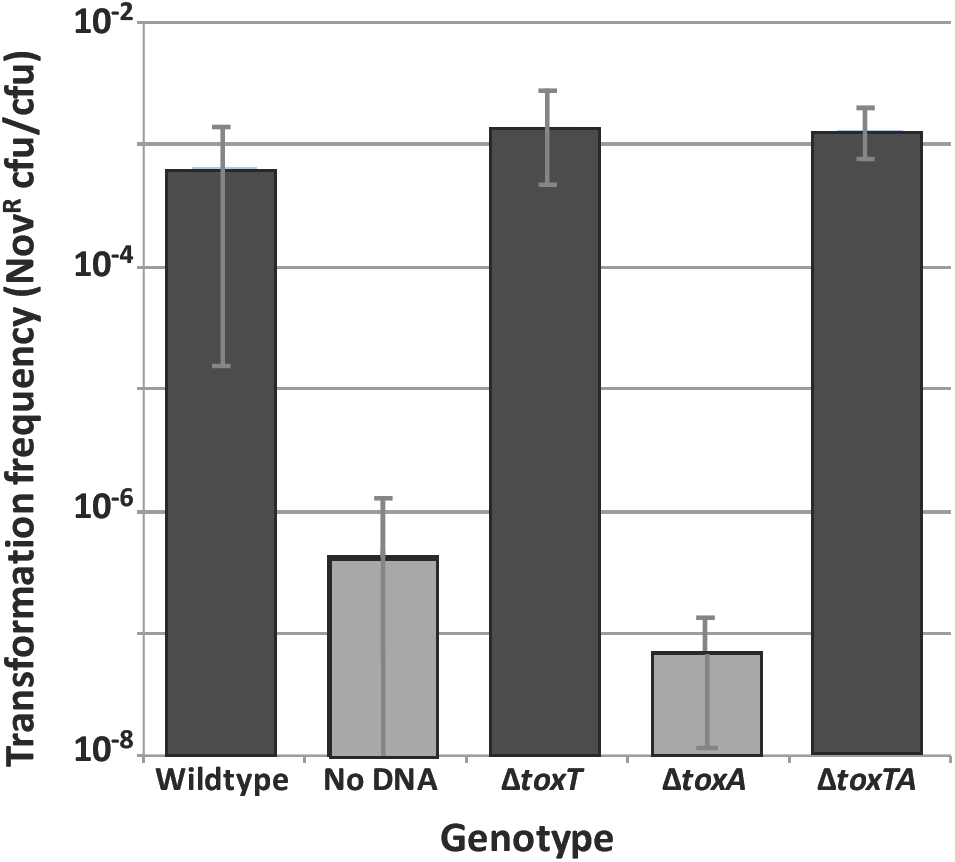
Transformation phenotypes of wildtype cells and *toxTA* mutants. Bars represent the means of at least three biological replicates, with error bars representing one standard deviation. Grey bars indicate values below the detection limit.

### ToxT does not modulate growth and/or competence development in normal cells

The ToxT protein must have a strong effect on competent cells, since earlier work found that a *toxA* knockout strain had no detectable DNA uptake or transformation under standard competence-inducing conditions (Sinha *et al.* 2012, see also **Fig. 2**). DNA uptake by the *toxA* mutant was below the limit of detection (100-fold reduction), but the >10^6^-fold reduction in transformation frequency provided a more sensitive measure of the magnitude of the defect. A simple explanation for this phenotype would be that unopposed ToxT prevents competence by killing or otherwise inactivating the cells in which it is expressed; however, growth rates were very similar between wildtype and the three *toxTA* mutant strains growing in rich medium (**Supp. Fig. A**). However, because the *toxTA* promoter is regulated by a CRP-S site, its expression (and thus ToxT production) might be limited to competent cells even in the absence of ToxA. This prompted us to look for evidence of competence-dependent toxicity. Cells with and without *toxA* had similar CFU/ml values in standard transformation assays, but this is not a very sensitive test of competence-dependent toxicity since cells in the competence-inducing starvation medium MIV are already growth-arrested. Furthermore, only 10%-50% of cells in a competent culture are typically transformable (Mell *et al.,* 2014), and the extent of competence gene expression in the non-transformable fraction is not known.

A more detailed time course analysis compared wildtype and Δ*toxA* mutant cell numbers (CFU/ml; **Fig. 3**) and culture densities (OD_600_; **Supp. Fig. B**) during growth in rich medium, during the development of competence, and during recovery from MIV starvation medium in rich medium. Δ*toxA* cells grew slightly slower than wildtype during the initial log phase growth in rich medium. In both cultures, transfer to the starvation medium MIV slowed growth to the same extent (both OD and in CFU/ml) (grey-shaded area in both figures). If unopposed toxin expression during competence development kills cells or halts growth, then returning cells to rich medium might reveal a stronger growth defect. However, both strains also had similar recovery kinetics after a fraction of their culture was returned to rich medium, although Δ*toxA* cells again grew slightly slower than wildtype.

**Figure 3:**
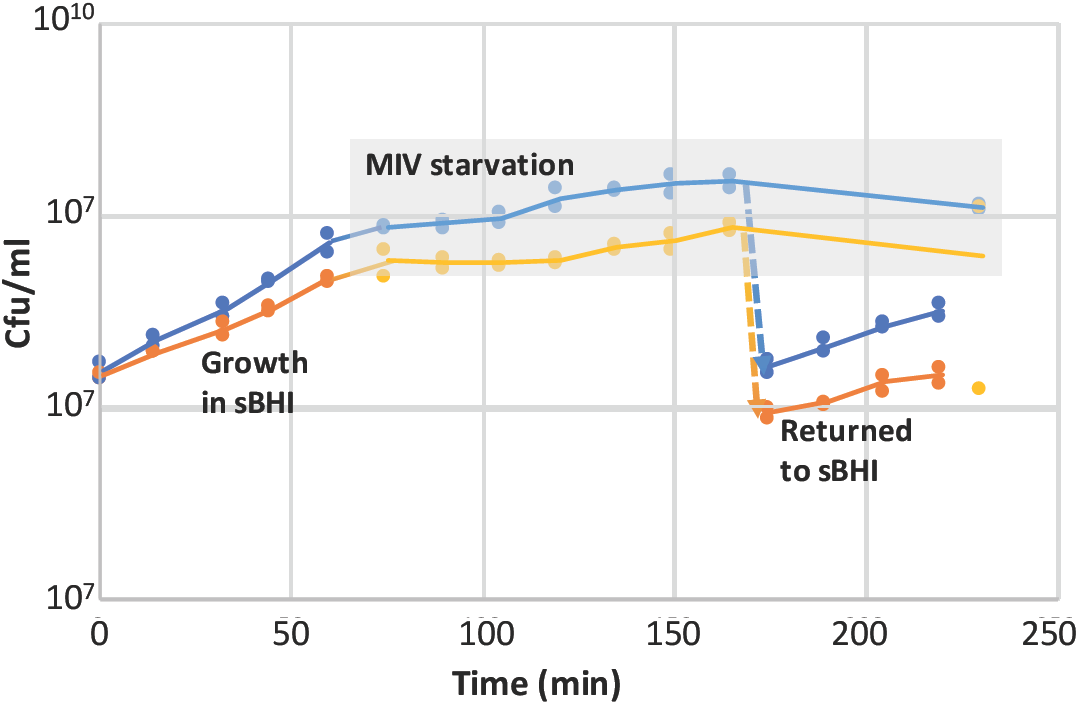
Growth and MIV recovery of log-phase KW20 and Δ*toxA.* Log-phase cells in sBHI were transferred to MIV at t=65 min; a portion of each MIV culture was diluted 10-fold into sBHI at t=170 min. The grey-shaded area indicates samples taken from MIV cultures. Blue: KW20, orange: Δ*toxA*.

Since cyclic AMP is required for induction of the competence genes, and addition of cAMP induces partial competence during exponential growth (Dorocicz *et al.,* 1993), we also tested the effect of cAMP on the Δ*toxA* knockout. Addition of cAMP did not rescue its transformation defect (Supp. Fig. C), so failure to transform is not caused by defective cAMP production in the antitoxin mutant.

Because chromosomal toxin-antitoxin systems are often reported to have acquired roles in modulating cell growth (Page and Peti, 2016), we examined the Δ*toxT* mutant for changes in growth and competence under various conditions. The grey line in Supplementary Fig. A shows that Δ*toxT’s* growth is indistinguishable from that of wildtype cells (blue line), and Fig. 2 shows that its MIV-induced competence is also unchanged. Supp. Fig. D shows that the kinetics of competence development and loss during growth in rich medium is also indistinguishable from wildtype. Also unchanged is the gradual loss of competence when cells are left overnight in MIV medium (not shown). We conclude that ToxT’s normal expression in wildtype cells does not detectably regulate competence development or loss. Since these phenotypic analyses did not show direct evidence of MIV-specific toxicity, we used RNA-seq to investigate how a *toxA* deletion affects expression of *toxA* and *toxT*, and how these changes affect cells.

### Transcriptional control of competence

The *toxTA* operon has a typical CRP-S-type regulatory motif upstream of the *toxT* coding sequence, and previous global analysis of transcription using microarrays (Redfield *et al.,* 2005) showed that it is competence-induced. We have now investigated this regulation in more detail as part of a comprehensive RNA-seq analysis of competence-associated gene expression in wildtype and mutant cells. In these experiments, samples for RNA preparations were taken from three replicate cultures at four time points, first when cells were growing in log phase in the rich medium sBHI (t=0), and then at 10, 30 and 100 minutes after each culture had been transfered to the competence-inducing starvation medium MIV.

We first examined how competence induction changed expression of known CRP-regulated (CRP-N) and CRP+Sxy-regulated (CRP-S) genes. Fig. 4 gives an overview of the results. Each coloured dot represents a gene, colour-coded by function. Its horizontal position indicates its level of expression in rich medium (T=0) and its vertical position indicates how this expression changed at later time points (**A**: T=10; **B**: T=30; **C**: T=100) or in a mutant background at T=30 vs. T=0 (**D**; Δcrp; **E**: Δsxy). Thus in Fig. 4A the higher positions of the green dots (genes regulated by CRP-N sites) and the red diamond (the competence regulator *sxy*) indicate that they were strongly induced after 10 min in MIV. Induction of *sxy* was followed (T=30 and T=100) by strong induction of the known competence-regulon genes (higher positions of CRP-S genes; blue dots) (Fig. 4**B** and **C**). Consistent with prior studies (Redfield *et al.,* 2005), induction of all these genes was blocked by deletion of the *crp* gene (Fig. 4**D**), and induction of the competence regulon (CRP-S) genes was blocked by deletion of *sxy* (Fig. 4**E**).

**Figure 4:**
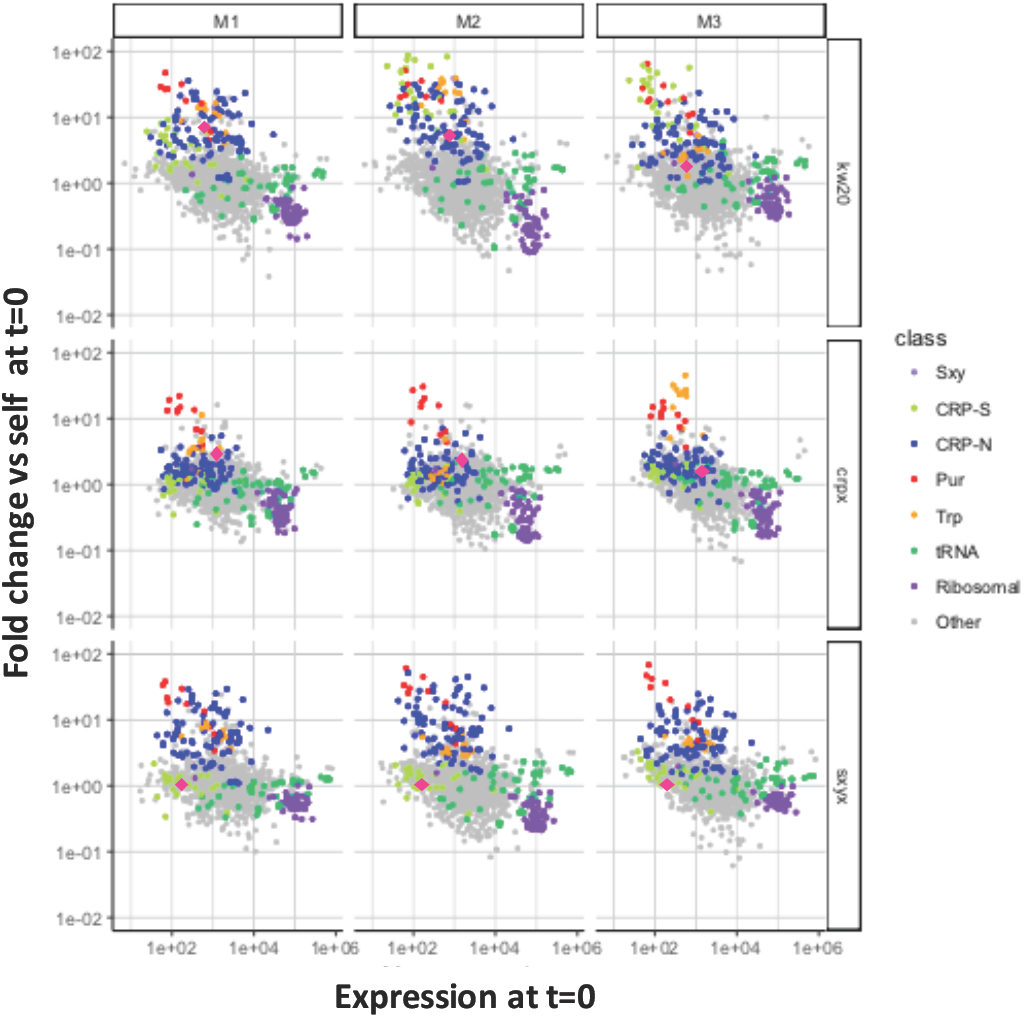
Changes in expression of genes regulated by CRP and Sxy. Each circle represents a gene, colour-coded by function: red: *sxy*, green, CRP-N-regulated; blue, CRP-S-regulated; yellow, ribosomal; purple, purine synthesis; orange, tryptophan synthesis; grey, other or unknown function. Each circle’s horizontal position indicates the gene’s level of expression in rich medium (T=0) and its vertical position indicates how this expression changed at later time points (**A**: T=10; **B**: T=30; **C**: T=100) or in a mutant background at T=30 (**D**; Δ*crp*; **E**: Δ*sxy*).

To identify all genes whose wildtype expression differed between sBHI and MIV, DESeq2 was used to compare RNA-seq expression values in wild-type RNA samples before and after competence induction (T=0 and T=30). Of the 1747 genes examined, 325 had significantly different expression (adjusted p-value < 0.05 after performing a Wald test and Benjamini-Hoschberg correction in DESeq2). To focus on genes with large changes in expression, we imposed an additional requirement that expression be changed by at least 4-fold, the same threshold used in the previous microarray study (Redfield *et al., 2005)*. This higher stringency gave 123 genes significantly decreased and 71 significantly increased, for a total of 194/1747 or 11% of all genes tested (listed in Supp. Table 2). Of these, 130 were among the 192 genes previously identified as differentially expressed in the microarray study.

Many of these changes are likely due to the absence of many essential nutrients from MIV medium. The significant induction of 9 of the 10 genes regulated by PurR is expected, since MIV lacks nucleotide precursors; the tenth PurR-regulated gene, HI1616, was also strongly induced in all replicates. MIV also lacks tryptophan, and all 11 Trp-regulon genes regulated by TrpR were significantly induced. The nutritional downshift is also likely to be responsible for the induction of many permeases and transporters, and for the significant downregulation of 39 of the 51 genes encoding ribosomal proteins (*rps* and *rpl* genes). The other 22 ribosomal protein genes were also downregulated, the majority with adjusted p-values < 0.05. Expression of 16S and 23S rRNAs was not measured since these molecules had been depleted from the samples during sequencing library prepation, but expression of the tRNA^Ala^, tRNA^Leu^, and tRNA^Gly^ genes encoded within the six rRNA operons was also significantly reduced.

To clarify the roles of competence-regulating genes in the observed MIV-induced gene expression changes, RNA-seq coverage values from *sxy* and *crp* knockout strains were compared to values for KW20 sampled at the same four sBHI and MIV timepoints. To evaluate changes at all timepoints simultaneously, a likelihood ratio test was performed using DESeq2 to identify genes that behaved differently between strains. Significant genes were flagged after adjusting for multiple hypothesis testing. Sxy is the competence-specific regulator, and deleting it significantly reduced expression of 24 of the known competence genes (Redfield *et al.* 2005, Sinha *et al.,* 2013). Two other previously identified competence genes, HI0250 (*ssb*) and HI1631, also had reduced expression, but these did meet the significance cutoff. *ssb* is an essential gene (Sinha *et al.,* 2012); its high baseline expression increased 40% on transfer to MIV and returned to normal in the *sxy* knockout. Expression of HI1631 was reduced 6-fold by the *sxy* knockout, but high variability in KW20 expression led to an insignificant adjusted P-value.

### Transcriptional control of *toxTA*

RNA-seq analysis confirmed that *toxTA* is regulated as a typical competence operon. In wildtype cells, baseline RNA-seq expression of *toxT* and *toxA* was very low during log phase in rich medium, with approximately tenfold induction by incubation in MIV (Fig. 5 (*toxT*) and Supp. Fig. E (*toxA*), green lines and points). As expected, this increase was eliminated by knockouts of CRP and Sxy (brown and blue lines and points). Like other CRP-S genes, both *toxT* and *toxA* were also induced in rich medium in the presence of mutations known to cause hypercompetence (Redfield 2005) (RNA-seq data not shown). Thus the *toxTA* operon is regulated as a typical member of the competence regulon.

**Figure 5:**
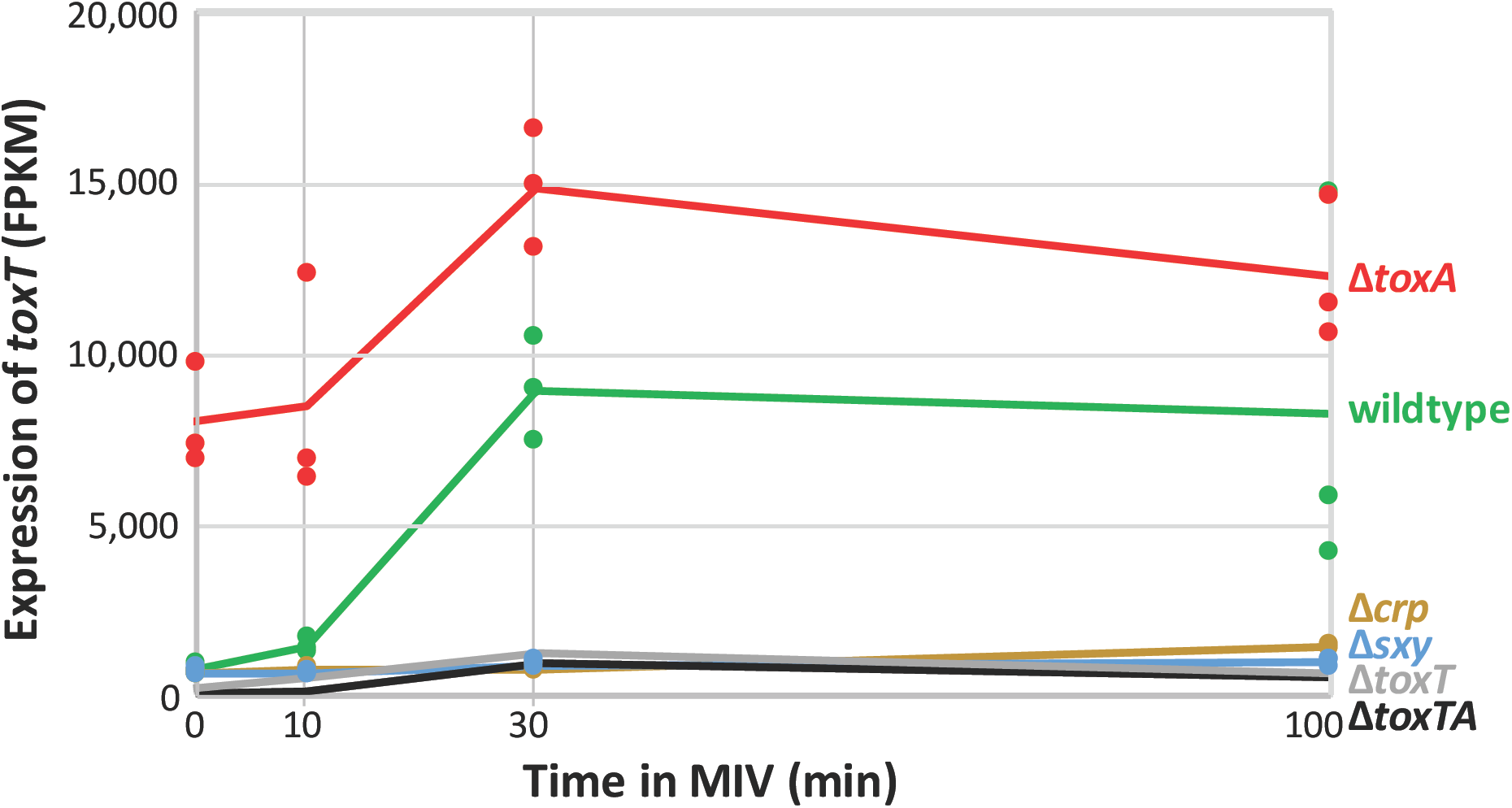
Competence-induced expression of *toxT*. Sample FPKM values (dots) and means (lines) for *toxT* (HI0660). Strains: wildtype: green; Δ*crp*: brown; Δ*sxy*: blue; Δ*toxA*: red; Δ*toxT*: grey; Δ*toxTA*: black. The values for the Δ*toxT* and Δ*toxTA* samples are underestimates because most of the gene has been deleted in these strains.

RNA-seq analysis also showed that the *toxTA* operon is regulated as a typical type II toxin-antitoxin operon. In such operons, the antitoxin protein usually protects cells from the toxin in two ways. First, it inactivates the toxin protein by forming a complex with it that has no toxin activity. Second, this toxin-antitoxin complex binds to the *toxTA* promoter and represses transcription (Overgaard, 2008, Goeders and Van Melderen, 2014). ToxA has a HTH-XRE DNA-binding domain, which is commonly found in promoter-binding antitoxins (Makarova *et al*., 2009, Yamaguchi *et al.,* 2011), and the RNA-seq analysis in Fig. 5 confirmed that it represses *toxTA* transcription. The Δ*toxA* mutant, which retains an intact *toxTA* promoter and *toxT* coding sequence (see Fig. 1), had 9-fold increased baseline expression of *toxT* in log phase cells (red line and points in Fig. 5). Expression increased further during competence development, with the same kinetics as in wildtype cells, suggesting independent contributions from baseline repression by antitoxin and competence induction by CRP and Sxy. (Values for *toxA* expression are shown by the red points and line in Supp Fig. E, but are underestimates because most of the gene has been deleted.)

Since antitoxin is predicted to repress *toxTA* only when bound to toxin, we were initially surprised that knocking out *toxT* or both *toxT* and *toxA* did not increase RNA-seq coverage of *toxT* (Fig. 5, grey and black lines) and that knocking out both genes did not increase coverage of *toxA* (grey line in Supp Fig. E). These mutants retain all the upstream sequences and the *toxT* start codon, and enough sequence of the deleted genes to identify them in the RNA-seq analysis.) An explanation was suggested by a recent study of the *Escherichia coli hicAB* toxin/antitoxin system (Turnbull and Gerdes, 2017), and confirmed by more detailed analysis of *toxTA* transcripts. The HicA (toxin) and HicB (antitoxin) proteins have no detectable sequence homology to ToxT and ToxA, but their operon is also Sxy-regulated and has the same atypical organization (toxin before antitoxin) (Sinha 2009). Turnbull and Gerdes show that the *hicAB* operon has two promoters. Promoter P1 has a CRP-S site regulated by CRP and Sxy, which is not repressed by the HicB antitoxin. A secondary promoter P2 is very close to the *hicA* start codon; it is repressed by HicB independently of HicA, and its shortened transcripts produce only functional HicB, not HicA. Promoter P1 of this *hicAB* system thus resembles the CRP-S regulation of the *toxTA* operon, and the presence of a second antitoxin-regulated internal promoter similar to P2 would explain the high *toxTA* operon expression seen in the *toxA* knockouts.

This finding in the *hicAB* system prompted us to do a more detailed analysis of *toxTA* transcription patterns in wildtype and mutant cells to determine whether the *toxTA* transcripts expressed in the absence of *toxA* were similarly truncated. Figure 6A shows RNA-seq coverage of the *toxTA* promoter region and the 5’ half of *toxT,* in wildtype cells and in the *toxA* deletion mutant (note that transcription of *toxTA* is from right to left). As expected, the predicted CRP-S promoter upstream of *toxTA* was active only at T=30 and T=100; its activity was not affected by deletion of *toxA*. Deletion of *toxA* instead caused strong constitutive transcription from a second promoter (‘P2’), with reads beginning about 30 bp downstream of the *toxT* start codon. Transcripts produced from this start point are unlikely to produce active ToxT; the only other in-frame AUG in *toxT* is 30 bp from the end of the gene, and it and the first GUG (position 35) lack Shine-Dalgarno sequences. This supports the hypothesis that the *H. influenzae toxTA* operon is regulated similarly to the *E. coli hicAB* operon, with the antitoxin repressing transcription from a downstream ‘P2’ promoter whose transcript produces antitoxin but not toxin.

**Figure 6:**
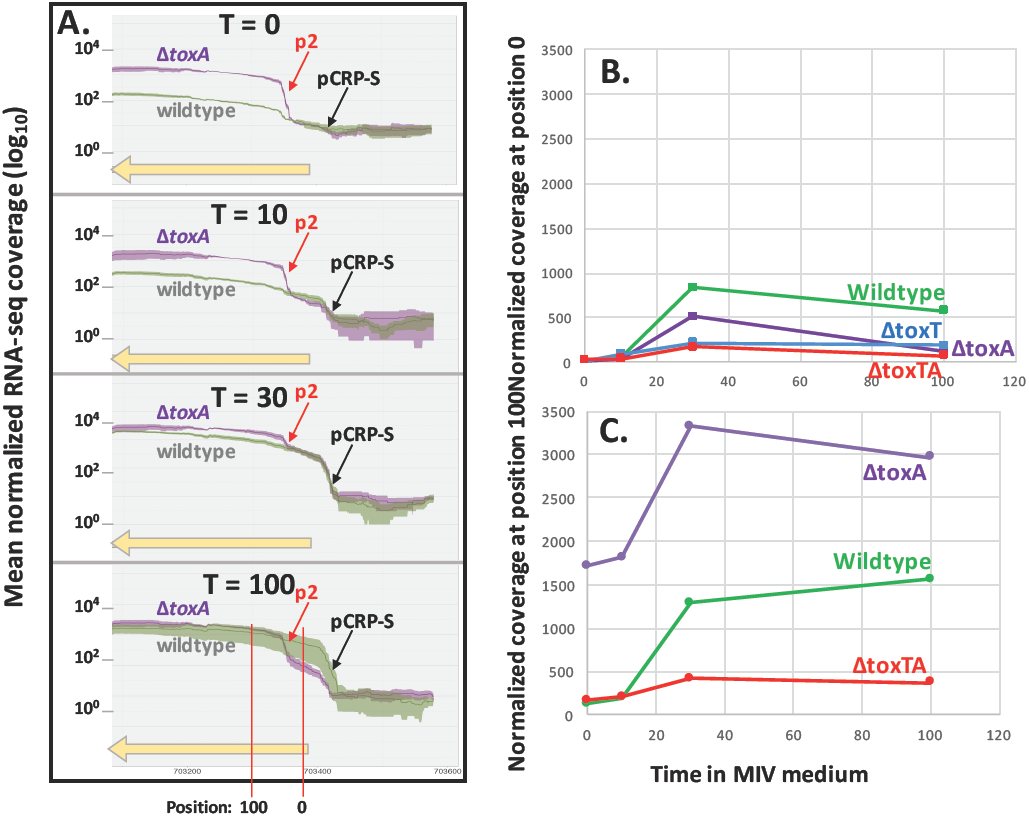
Read coverage of the toxTA promoter region. **A.** The green (KW20) and purple (ΔtoxA) lines indicate mean normalized coverage at each position, shaded areas indicate standard errors. The yellow bar indicates the 5’ half of *toxT*. **B.** and **C.** Time course of normalized read coverage at two specific positions in the toxTA operon. **B.** Position 0 = *toxA* start codon. **C.** Position 100.

In the *E. coli hicAB* system, P2 is repressed by HicB antitoxin alone, binding of HicB to the P2 operator is destabilized when HicA toxin is abundant, and transcription from P2 in plasmid constructs is elevated when the chromosomal *hicAB* operon is deleted (Turnbull and Gerdes, 2017). To see if this also happens in *H. influenzae*’s *toxTA*, we measured transcription in wildtype and *toxTA* mutant cells more accurately by scoring the coverage at two positions in the *toxTA* operon (indicated by red vertical lines at the bottom of Fig. 6A). Position 0 is the *toxT* start codon, 34 nt downstream from the CRP-S promoter (P_CRP-S_) but upstream of the putative P2 promoter, and position 100 is about 70 nt downstream from P2 (P2 and position 100 are deleted in Δ*toxT*). To eliminate read-mapping artefacts arising from failure to align reads that span an insertion or deletion, each mutant’s reads were mapped onto its own *toxTA* sequence rather than the reference sequence. Comparison of Figures 6B and 6C shows that coverage at position 100 was always higher than coverage at position 0, consistent with the presence of a second promoter between positions 0 and 100. Fig. 6B also shows that coverage at position 0 (expression from P_CRP-S_) was reduced by all of the *toxTA* deletions. This was unexpected, and suggests that this promoter may have unusual properties, since coverage of other CRP-S genes was not similarly affected. The *toxA* deletion caused the predicted increase in coverage at position 100 (Fig. 6C), but the *toxTA* deletion unexpectedly reduced rather than increased coverage at this position ∼3-fold from the wildtype level, even though this construct retains the first 150 bp of the operon, including P2. This reduction was not accounted for by the reduction in expression from P_CRP-S_, suggesting that high-level transcription from the *toxTA* P2 promoter only occurs when ToxT is present and ToxA is absent. This could mean either that ToxT directly binds the P2 promoter to induce transcription, which seems unlikely given its lack of DNA-binding domain, or it could mean that the presence of ToxT disrupts binding of a secondary repressor of the operon, such as a noncognate antitoxin (Goeders and Van Melderen, 2014).

### ToxT does not prevent induction of the competence regulon

To investigate how deletion of the *toxTA* antitoxin causes severe defects in DNA uptake and transformation, we first examined changes in expression of the genes that regulate the competence regulon. Comparison of the RNA-seq data for wildtype cells (orange) and *toxT, toxA* and *toxTA* mutants (yellow, blue and grey) ruled out direct inhibition of competence gene expression by a *toxT*-encoded toxin. Unopposed expression of *toxT* (in the Δ*toxA* mutant) only slightly reduced induction of the *sxy* transcript needed for induction of the competence regulon (Supp. Fig. F-A, blue line). Importantly, similar modest reductions were also seen in the other *toxTA* mutants (grey and yellow lines), which have normal competence. The mRNA levels of *crp* and *cya* (CRP (HI0957) and adenylate cyclase (HI0604)) were also not changed by Δ*toxA* (Supp. Fig. F-B and F-C).

The competence operons induced by these regulators also retained normal or near-normal expression in the Δ*toxA* mutant at 30 min after transfer to MIV, the time when competence-induced gene expression is highest (Fig. 7). As noted above for *sxy*, competence gene expression levels at this time were very similar between the Δ*toxA* mutant, which cannot take up DNA or transform, and the Δ*toxT* and Δ*toxTA* mutants, which take up DNA and transform normally (Supp. Fig. G-A and G-B). Although expression of most of the competence operons was substantially reduced in Δ*toxA* cells at the t=100 min time point (Supp. Fig. G-C), this effect was too little and too late to explain the mutant cells’ complete lack of competence.

**Figure 7:**
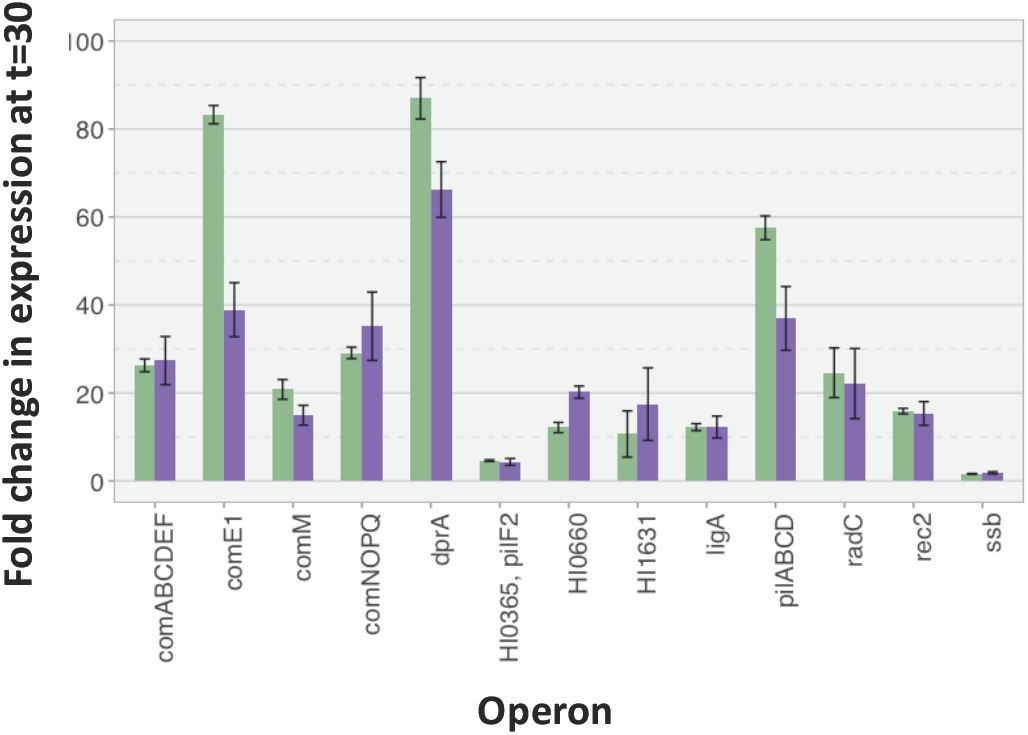
Changes in competence operon expression levels after 30min in MIV. Fold changes in competence operon expression levels in KW20 (green) and Δ*toxA* (purple) after 30 min in MIV, compared to 0 minute samples. Black lines show standard errors.

### Other ToxT and ToxA effects in competence-induced cells

Since changes in competence gene expression could not readily explain the severe competence defect of Δ*toxA*, we extended our investigation to genes not known to be involved in competence. Supp. Table 3 lists, for each timepoint, the genes whose expression was significantly different in the Δ*toxA* mutant than in all three strains with normal competence (wildtype, Δ*toxT* and Δ*toxTA*). In rich medium (T=0) the only statistically significant effect of Δ*toxA* on gene coverage was about 1.5-fold increased expression of three genes in the HI0654-0658 operon, which are directly downstream from *toxA* (see Fig. 1) and thus may experience read-through from the *toxTA* P2 promoter (which was constitutively active in the Δ*toxA* mutant).

The operon includes genes encoding shikimate dehydrogenase, an ABC transporter, and a hypothetical protein with putative topoisomerase I domains. Expression of genes in this operon increased about 1.2-1.5-fold in MIV in wildtype cells and in other mutants with normal competence, suggesting that read-through also occurs from the *toxTA* CRP-S promoter. Their normal induction in competence suggests that their higher expression in Δ*toxA* is unlikely to be responsible for this strain’s competence defect, but it may cause the slight Δ*toxA* growth defect described above (Supp. Fig. A). The absence of other detectable changes in gene expression is consistent with the postulated lack of functional ToxT protein produced from the P2 promoter.

ToxT effects are also not expected at the 10 min time point, since the Sxy-dependent CRP-S promoter is not yet active. Only two genes were significantly changed in Δ*toxA*: HI0655 (see above) and HI0231 (*deaD*), which encodes a DEADbox helicase involved in ribosome assembly and mRNA decay (Iost and Dreyfus, 2006). In all strains, this gene’s expression falls rapidly on transfer to MIV, but levels in Δ*toxA* were about 50% higher at all time points.

At the 30 min time point, seven genes’ expression levels were significantly altered by deletion of *toxA*. Most were only changed by about 2-fold, but two genes had large increases at both the t=30 and t=100 time points and may be relevant to the competence defect: Deletion of *toxA* increased HI0235 expression 3-5-fold at t=30 and 2-3-fold at t=100 (in the Δ*toxT* and Δ*toxTA* comparisons, but not in the KW20 comparison. Its protein has strong similarity to the ArfA ribosome-rescue domain (Garza-Sanchez *et al*., 2011); the significance of this is discussed below. HI0362 encodes a CRP-regulated iron-transport protein that normally increases in MIV but does not increase in *toxA* deletion mutants.

Global RNA-seq analysis did not reveal any obvious candidate genes. Although many more genes were significantly changed by Δ*toxA* at the 100 min time point, only four of these were also changed at t=30. Two of these, HI0235 and HI0362, were described above. Additionally, In all competent strains, HI0504 (*rbsB*, a ribose transporter component), was induced 20-fold more in MIV than other genes in its operon, but this increase was only 10-fold in Δ*toxA* (at t=30 as well as t=100). Expression of HI0595 (*arcC*, carbamate kinase) normally falls 2-3-fold immediately after transfer to MIV, but the fall was greater in Δ*toxA*. 28 other genes were significantly changed only at t=100, but their expression patterns and predicted functions were diverse and did not suggest an explanation for Δ*toxA*’s lack of competence.

Overall, this gene expression analysis did not reveal any promising mechanisms through which unopposed *toxT* expression could prevent competence.

### Related toxins may suggest mechanism of action

Since examination of gene expression shed little light on how the ToxT toxin prevents competence, as an alternative approach we considered the modes of action of well-studied relatives of ToxT. The most common type II toxins act as translation-blocking ribonucleases, such as RelE, but several alternative modes of action are also known, and some newly discovered toxins lack identified activities (Makarova *et al.,* 2009). The Pfam and TAfinder databases assign the *H. influenzae* ToxT protein to the ParE/RelE toxin superfamily, whose characterized members include both ribonucleases and gyrase inhibitors (Goeders and Van Melderen, 2014). Because *toxTA* shares regulatory features, gene order, and chromosomal location with *E. coli’s hicAB*, we gave special consideration to the possibility that their toxins also share a mechanism; the HicA toxin is a ribonuclease that arrests cell growth by cleaving mRNAs and other RNAs (Jorgensen et al., 2009).

### Unopposed toxin does not inhibit gyrase

If ToxT inhibited gyrase we would expect the RNA-seq data to show that transfer to MIV caused increased expression of *gyrA* (HI1264) and *gyrB* (HI0567) and reduced expression of *topA* (HI1365), since these genes have opposing activities and compensatory regulation by DNA supercoiling (Gmuender *et al*., 2001). However, these genes’ coverage levels were similar in wildtype and all *toxTA* mutants, during both exponential growth and competencedevelopment.

### Unopposed toxin does not cleave competence-induced mRNAs site-specifically

The best-studied homologs of the *toxT* toxin act by cleaving mRNAs at random positions near their 5’ ends during their translation on the ribosome (Hurley, 2011, Goeders *et al.,* 2013). Thus we considered whether ToxT might prevent competence by one of two mechanisms. First, ToxT might specifically cleave the 5’ ends of competence-gene transcripts, eliminating their function without significantly changing their overall RNA-seq coverage levels or otherwise interfering with essential cell functions. Visual examination of RNA-seq coverage of all positions within the competence operons did not reveal any anomalies that might indicate that the mRNA in Δ*toxA* cells had been inactivated either by cleavage at specific sites or by random cleavage near the 5’ end (Gordon *et al.,* 2017). As an example, Supp Fig. H compares read coverage across the *comNOPQ* operon in wildtype and Δ*toxA* cultures after 30 min in MIV.

### Unopposed toxin may nonspecifically cleave mRNAs

A second mechanism we considered was that ToxT might nonspecifically cleave mRNAs. This would result in a large population of mRNAs lacking in-frame stop codons (‘non-stop’ mRNAs). Because these cannot undergo the normal ribosome-release process, this would cause a general block to translation (Tollervey, 2006). This block in turn is predicted to arrest cell growth until normal translation can be restored (Pandey and Gerdes, 2005). To indirectly detect such cleavage, we examined the insert sizes of our RNA-seq sequencing libraries by comparing the spanning length distributions of paired-end sequencing reads among strains.

Because independent library preparations had different insert sizes, comparisons were limited to samples prepared at the same time. Fig. 8 shows that the Δ*toxA* samples from library batch 1 had shorter fragment sizes than the KW20 samples from the same batch, and that the difference increased as the time after competence induction increased. This supports the hypothesis that the extreme lack of competence in Δ*toxA* cultures is due to non-specific ToxT cleavage of mRNAs.

**Figure 8:**
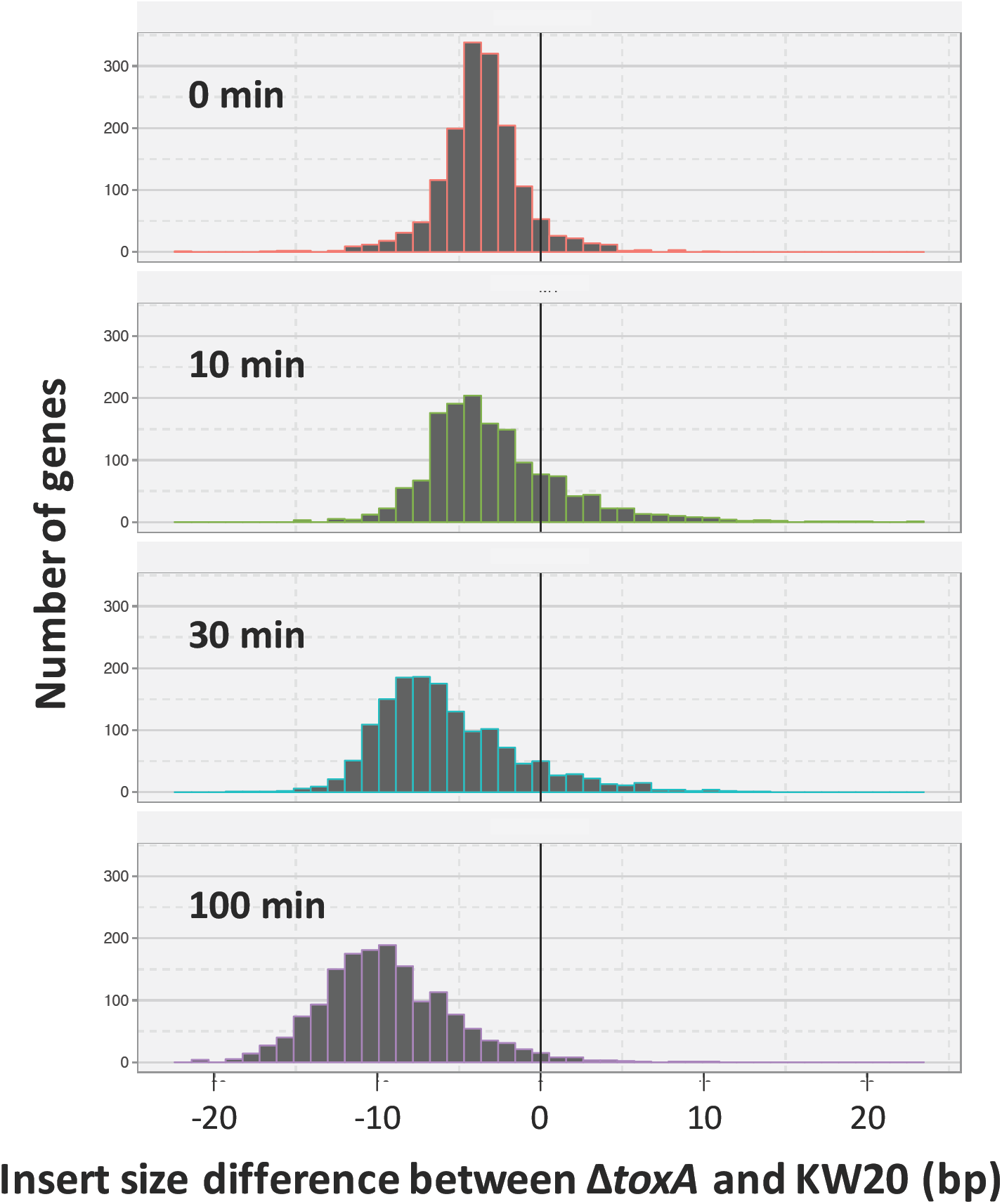
Distribution of insert-size differences between RNA-seq libraries prepared at the same time. Distribution of insert length differences between KW20 (kw20_A and kw20_B samples) and Δ*toxA* (antx_A samples) after 0, 10, 30 and 100 minutes in MIV.

Additional support for a generally toxic effect on translation comes from the *toxA* deletion’s effects on genes known to rescue ribosomes that have stalled on non-stop mRNAs. *H. influenzae* has two rescue systems: In the first, an abundant small RNA named transfer-messenger RNA (tmRNA), binds with its protein cofactor SmbB to arrested ribosomes, detaches both the non-stop mRNA and the incomplete protein, and tags the protein for degradation. In the second rescue system, ArfA recruits ribosome release factor 2 (HI1212) to the ribosome and causes it to cleave the nascent peptidyl-tRNA (Keiler, 2015). Translation of *arfA* is increased when tmRNA activity is reduced (Garza-Sanchez *et al.,* 2011). Consistent with these expectations, tmRNA (HI1281.2) is downregulated in the Δ*toxA* mutant, however, as noted above, the *arfA* homolog HI0235 is upregulated several fold (Christensen et al. 2003a, 2003b).

### Sxy regulation of TA systems

Jaskolska and Gerdes (2015) and Sinha *et al.,* (2009) reported that three other *E.* coli TA operons are regulated by Sxy, so we examined the promoter sequences and expression levels of the other seven *H. influenzae* TA operons. None of the promoters had strong matches to the *H. influenzae* CRP-S consensus (Cameron and Redfield, 2008, Sinha *et al.,* 2009) and their RNA abundance levels showed no evidence of competence-regulated expression or dependence on Sxy.

### Phylogenetic evidence for lateral transfer of the *toxTA*

Since toxin/antitoxin operons are highly mobile (Makarova *et al., 2009)*, we examined the distribution of the *toxTA* operon in other strains and species (Fig. 9). *toxTA* operons are present at the same genomic location in most *H. influenzae* genomes (see below) and in the closely related *H. haemolyticus*, but there are no recognizable homologs in most other bacteria (including most other members of the Pasteurellaceae). Instead, most identifiable homologs (with about 60% identity) are in a very distant group, the Firmicutes, especially *Streptococcus* (96 of the top 100 BLAST hits to ToxT outside the Pasteurellaceae are to diverse *Streptococcus* species). This suggests that the *toxTA* operon may have been transferred from a Firmicute into a recent ancestor of *H. influenzae* and *H. haemolyticus*. When we excluded *Streptococcus* spp. from the BLAST search, sporadic matches were found in a wide variety of other taxa. In addition, *toxTA* operons with about 50% identity were found in one other small Pasteurellacean clade (*Actinobacillus sensu stricto*), and on two 11kb plasmids (pRGRH1858 and pRGFK1025) from an uncultured member of a rat gut microbiome and an uncultivated *Selenomonas* sp. The distribution is summarized in Fig. 9A.

**Figure 9:**
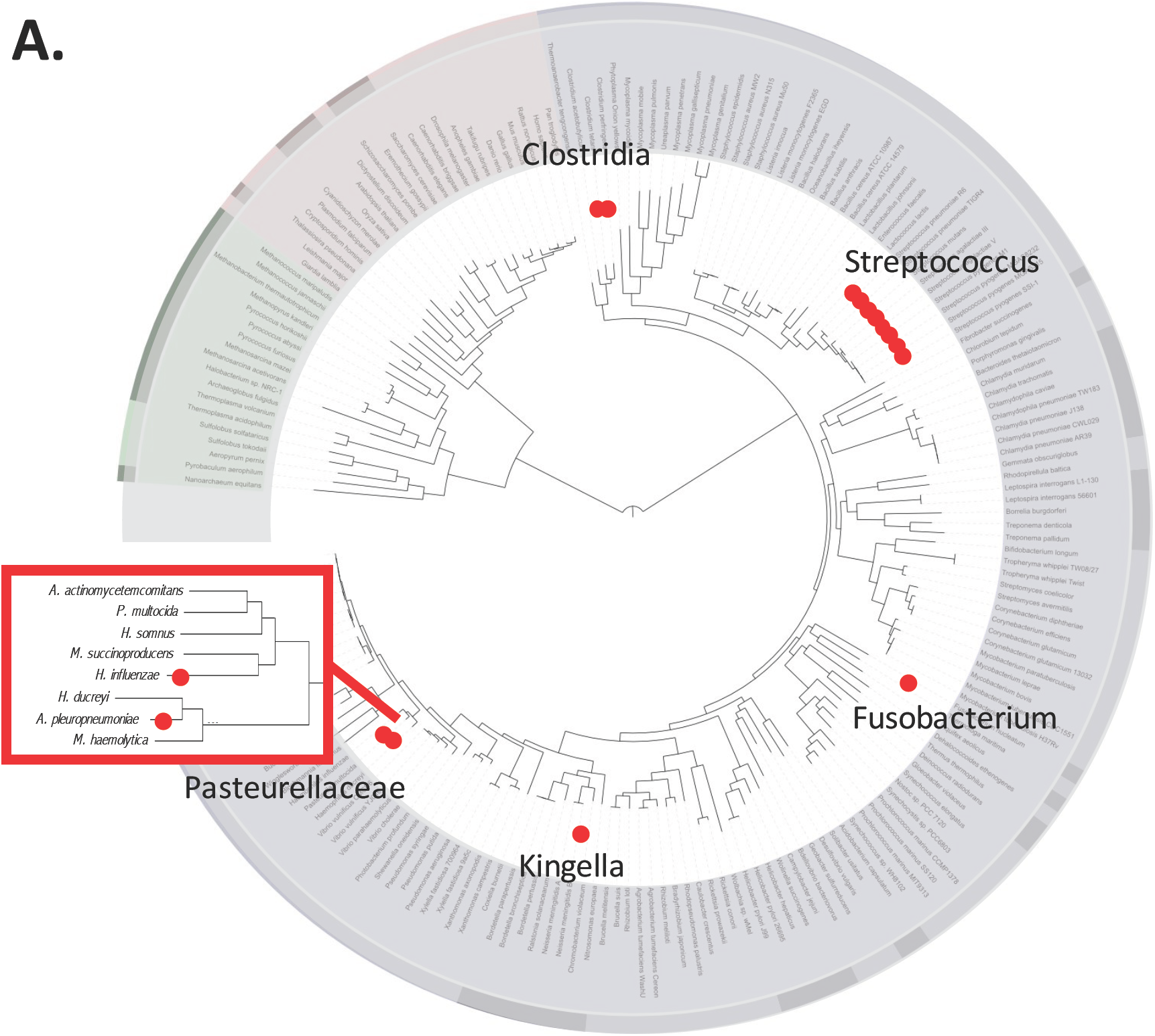
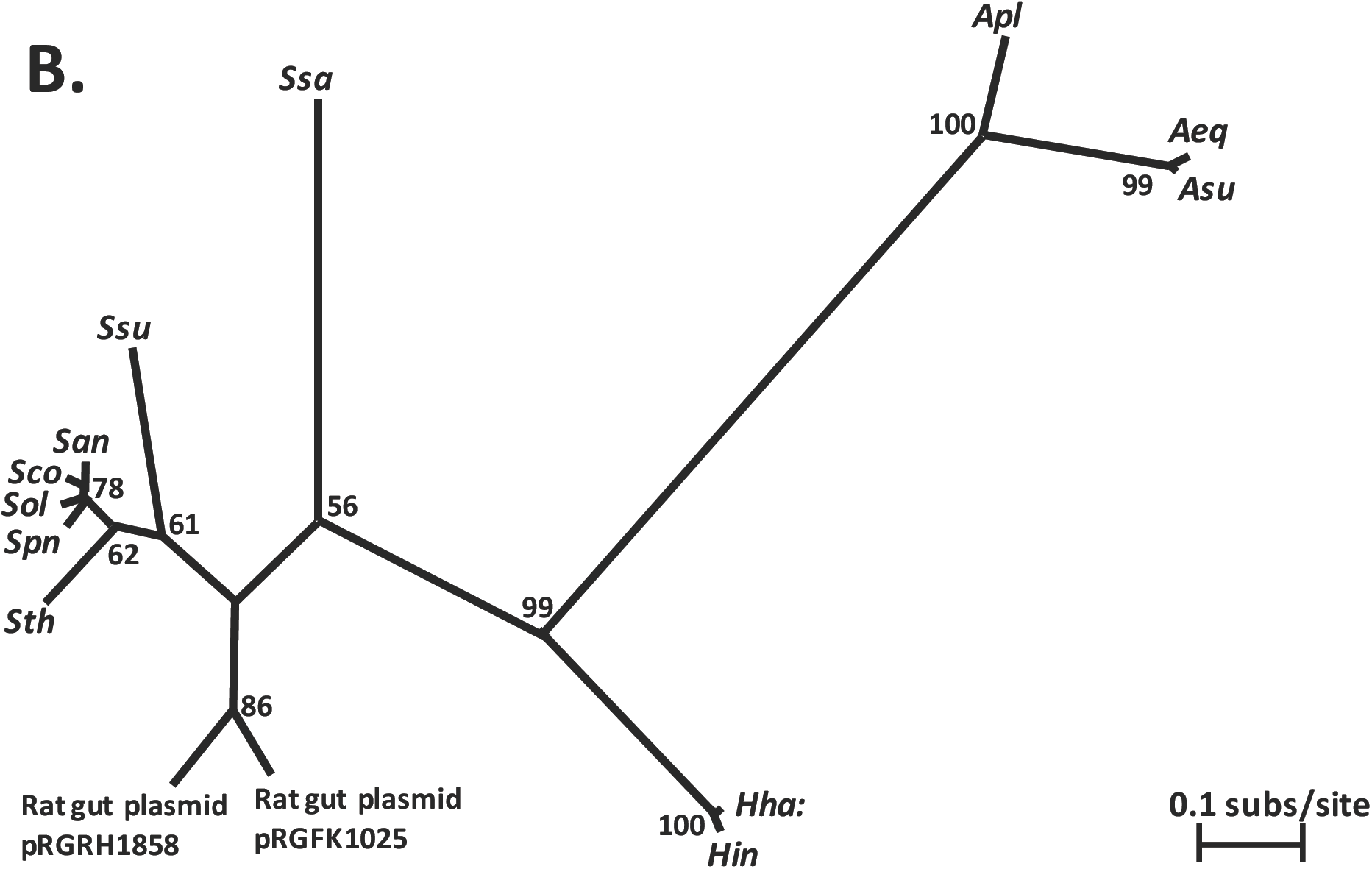
Distribution of *toxTA* homologs in bacterial genomes. **A.** Red dots indicate one or more taxa containing homologs of both ToxT and ToxA. Bacterial phylogeny image from Wikimedia Commons (Letunic 2007). Inset: Pasteurellacean phylogeny from Redfield et al. 2006. **B.** Unrooted maximum likelihood phylogeny of concatenated toxT and toxA homologs from selected species where both are present. Numbers at nodes are bootstrap values. Species abbreviations: *Apl: Actinobacillus pleuropneumoniae; Aeq: A. equuli; Asu: A. suis; Haemophilus haemolyticus; Hin: H. influenzae; Ssa: Streptococcus salivarius; Ssu: S. suis; San: S. anginosus; Sco: S. constellatus; Sol: S. oligofermentans; Spn: S. pneumoniae; Sth: S. thermophilus.*

To resolve the history of gene transfer events in the two Pasteurellaceae sub-clades, we created an unrooted maximum likelihood phylogeny of concatenated *toxT* and *toxA* homologs from selected species where both genes are present (Fig. 9B). Although there is 99% bootstrap support for a *Haemophilus*-*Actinobacillus* clade, the absence of homologs from all other Pasteurellaceae makes a single Pasteurellacean origin unlikely, since it would require there to have been multiple deletions in other Pasteurellacean subclades, or a second lateral transfer. Since the *Actinobacillus* sequences are also more distant from the *Haemophilus* sequences than from the *Streptococcus* sequences, the two Pasteurallacean groups may instead have acquired their *toxTA* operons by independent lateral transfers, probably from Firmicutes, since these homologs have the highest identity to the Pasteruellacean sequences. The alternative hypothesis of a single Pasteurellacean origin requires that acquisition was followed by multiple deletions, though the analysis in the next paragraph makes this less implausible.

### Deletions in *H. influenzae toxT* are common

181 *H. influenzae* genome sequences were available for examination. (Supp table #???) Of these, 162 had recognizable *toxA* sequences. All of these encoded full length ToxA proteins, but all except 24 had one of two common deletions affecting *toxT*. The extents of these deletions are shown by the dark grey bars at the bottom of Fig. 10. The most common deletion (n=93) removed 178 bp of *toxT* coding sequence but left both promoters intact. The second (n=45) removed 306 bp of sequence including both *toxTA* promoters and the *toxT* start codon. The 19 genomes that lacked recognizable *toxA* sequences all had the same 1015 bp deletion removing both *toxT* and *toxA* but leaving the flanking genes intact. In place of the missing sequences were 87 bp that have no homologs in GenBank. The average pairwise distance among the 162 *toxA* genes is 0.106, which is slightly higher than 0.088, the average of all genes with at most one copy per strain. The d_N_/d_S_ ratio of 0.037 is consistent with mild purifying selection on *toxA* and is higher than the average gene, which is 0.243. However, the strength of selection may be underestimated, since most *toxA*s lack functional *toxT* and/or may not be expressed due to deletions. Both sequence divergence and the high frequency of *toxT* deletions agree with expectations for a toxin/antitoxin system whose antitoxin protects against a toxin that is at least mildly deleterious.

**Figure 10:**
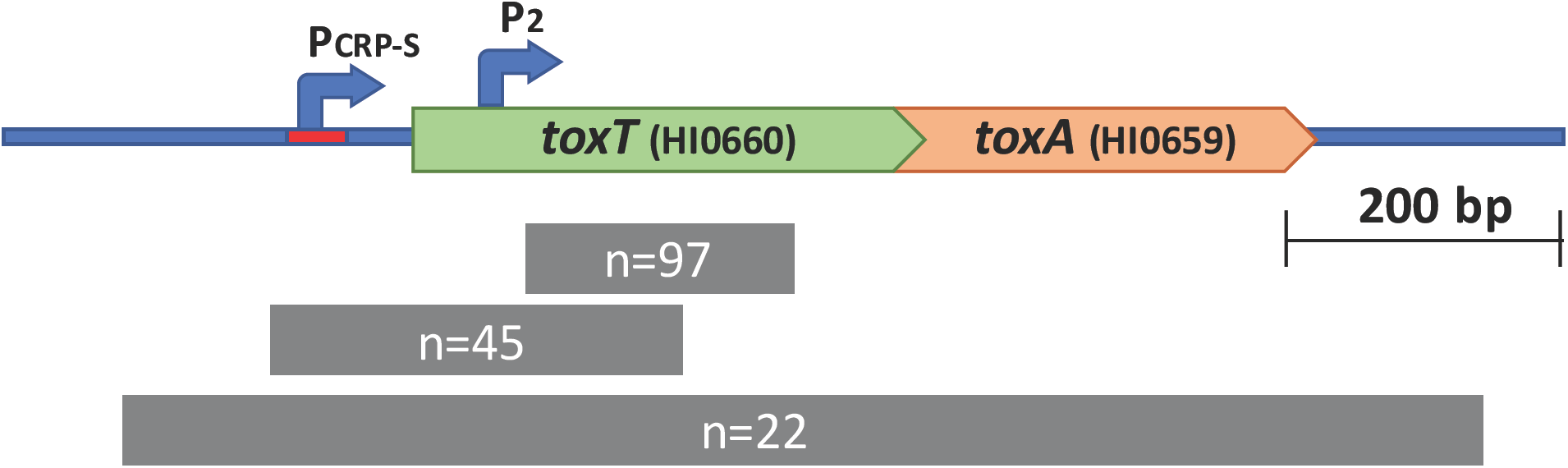
Natural deletions in the toxTA operon. Top line: structure of the wildtype toxTA operon in strain KW20. Lower lines: dark grey bars indicate the spans of the three naturally occurring deletions, annotated with number of strains possessing each deletion.

As mentioned above, the *E. coli* HicA and HicB protein sequences have little sequence similarity to ToxT and ToxA, and our wildtype *H. influenzae* strain (KW20, Rd) lacks a *hicAB* operon, but Syed and Gilsdorf (2007) found that 69/79 other *H. influenzae* strains were positive for *hicAB* by dot-blot analysis, so we examined the *hicAB* genes in our set of 181 *H. influenzae* genomes. Like *toxTA*, the *hicAB* operons in most *H. influenzae* strains have intact antitoxin (*hicB*) genes but deletions in their toxin (*hicA*) genes. Of the 181 strains examined, 122 were tagged as having *hicB*. All but 20 of these have a 250 bp deletion that removes both *hicAB* promoters and the first 50 bp of *hicA*. Many strains that lack *hicAB* share a large deletion that removes a large multi-gene 7147 bp segment flanked by a 57 bp duplication, but others have more complex structures that were not investigated further. Overall, the deletion pattern of the *H. influenzae hicAB* genes resembles that of *toxTA*, with frequent deletions of the toxin gene but preservation of the antitoxin.

### Might the variation in *toxTA* help explain the observed strain-specific variations in DNA uptake and transformation?

Maughan and Redfield (2009) measured the ability of 34 *H. influenzae* strains to both take up DNA and become transformed, so we examined this data for correlations with the presence of *toxTA* in the 19 of these strains whose *toxTA* and *hicAB* genotypes we were able to determine. All but one of the 19 strains had a complete *toxA* coding sequence but only five had intact toxTA operons. Of the rest, four had the large deletion that removed both *toxTA* promoters, nine had the smaller deletion internal to *toxT*, and one had the 1015 bp complete deletion. There was no obvious correlation between the *toxTA* genotypes and the DNA uptake or transformation phenotypes, but there was insufficient data for a high powered analysis.

### Does the *Actinobacillus pleuropneumoniae toxTA* operon affect competence?

The *A. pleuropneumoniae toxTA* operon was originally reported to have the CRP-S promoter typical of competence operons (Bosse *et al.,* 2009). Although reexamination of the promoter region failed to identify a high-quality CRP-S site, we constructed *toxT, toxA* and *toxTA* knockout mutants to investigate whether a *toxA* deletion would prevent competence. There were no significant differences between the transformation frequencies of wildtype cells and all *toxTA* mutants. Thus we conclude that the *A. pleuropneumoniae toxTA* operon does not affect competence. Expression of the *toxT* gene in the absence of the antitoxin had no detectable effect on growth or survival.

## DISCUSSION

Our investigation into why a HI0659 knockout prevents competence has provided a simple answer: HI0659 encodes an antitoxin (ToxA) needed to block the expression and competence-preventing activity of the toxin encoded by HI0660 (ToxT). But this answer has generated a number of new questions that we have only partially answered. Why is competence controlled by a toxin/antitoxin system? How does this system completely abolish DNA uptake and transformation without causing significant cell death? How did this TA system come to be competence-regulated? Does it confer any benefit to the cells, either generally or competence-specific?

Several findings support the conclusion that HI0660 and HI0659 encode proteins that function as a toxin/antitoxin pair. First is the similarity of the encoded ToxT and ToxA proteins to biochemically characterized toxin and antitoxin proteins of the RelE/ParE families. Second, and the strongest evidence, is the restoration of normal DNA uptake and transformation to antitoxin-knockout cells when the putative toxin is also knocked out. Third is the regulatory similarity between this system and the *hicAB* system of *E. coli*.

### How did the *toxTA* operon come to be in the *H. influenzae* genome and under competence regulation?

*H. influenzae* acquired its *toxTA* operon by horizontal transfer, either into a deep ancestor of the Pasteurellaceae or independently into more recent ancestors of *H. influenzae* and *A. pleuropneumoniae*. The closest relatives of the *toxTA* genes are in the distantly related Firmicutes, with homologs especially common in *Streptococcus* species. Since the Streptococci and Pasteurellaceae share both natural competence and respiratory-tract niches in many mammals, there may have been frequent opportunities for horizontal transfer between them.

We do not know how the *toxTA* operon came to be under CRP-S regulation. The *toxTA* operon’s strong regulatory parallels with the *E. coli hicAB* system suggest two explanations. One hypothesis is a distant shared evolutionary origin of the two systems, with selection maintaining regulation more strongly than protein sequence. Based on this hypothesis, the strong sequence similarity between the Pasteurellacean and Streptococcal *toxTA* systems then predicts that the regulatory features shared by the Pasteurellacean *toxTA* systems and the more-distant *hicAB* system (a competence-regulated promoter producing both proteins and an antitoxin-regulated promoter producing only antitoxin) could also be shared by the Streptococcal homologs. However, it is also possible that toxin-antitoxin systems with similar regulation and function have adopted similar roles in separate instances, a phenomenon which is more likely in toxin antitoxin systems as they undergo frequent horizontal transfers and are often under strong selective pressure. The *sxy* gene and the CRP-S promoters it regulates are not known outside of the Gamma-Proteobacteria sub-clade that contains the Vibrionaceae, Enterobacteraceae, Pasteurellaceae and Orbaceae (Cameron *et al.,* 2006). Thus, it would be interesting to examine the regulation and function of the *toxTA* homologs outside the Pasteurellaceae to determine when and where it adopted a regulatory role and the mechanism of the toxic activity. Examining these homologs could give insight into both the mechanism of action of the *H. influenzae toxTA* system, and its evolutionary history.

### How does unopposed ToxT prevent DNA uptake and transformation?

The transformation defect caused by deletion of the antitoxin gene toxA is very severe, so it was surprising that RNA-seq analysis detected only few and minor changes in expression of competence genes. Instead, the best explanation is that ToxT is an mRNA-cleaving ribonuclease, whose activity causes a general block to translation that prevents functioning of the induced competence genes. The most direct evidence is the decrease in insert size distributions seen in Δ*toxA* mutants, but this conclusion is also supported by the combination of regulatory similarities between the *toxTA* and *hicAB* systems and by sequence similarities between the ToxT protein and Type II ribonuclease toxins.

### Why then does the Δ*toxA* mutant not suffer from growth arrest or toxicity?

Part of the explanation is that mRNAs encoding functional ToxT are only expressed after cells have been transferred to competence-inducing starvation medium, a condition that severely slows cell growth and division even in wildtype cells. Detecting the predicted competence-specific toxicity is further complicated by the uneven distribution of transformability in competence-induced cells. Co-transformation experiments using multiple unlinked markers consistently show that no more than half, and sometimes as little as 10%, of the cells in a MIV-treated culture produce recombinants (Mell and Redfield, 2014). We do not know whether only the transforming cells express the competence genes or all cells express them but some fail to correctly assemble the DNA uptake or recombination machinery. If only a modest fraction of the cells in a competent culture are expressing the toxin then any toxic effect on culture growth and survival will be more difficult to detect.

### Does this operon confer any benefit (or harm) on *H. influenzae*?

Why have a competence-regulating toxin/antitoxin system at all, when it has no detectable effect on competence unless its antitoxin component is defective? Regulatory parallels with the *hicAB* system suggest that CRP-S regulation is not incidental. We found no direct evidence of any toxin-dependent alteration to the normal development of competence. Production of Sxy is subject to post-transcriptional regulation by the availability of nucleotide precursors (Macfadyen *et al*., 2001, Sinha *et al*., 2013), and we have elsewhere proposed that DNA uptake is an adaptation to obtain nucleotides when nucleotide scarcity threatens to arrest DNA replication forks (Mell and Redfield, 2014). In this context, competence-induction of the *toxTA* operon may be a specialization to help cells survive, by slowing or arresting protein synthesis until the nucleotide supply is restored.

However, the high frequency of deletions that remove either complete *toxTA* or both promoters (35%) indicates that the operon is dispensable. And the even higher frequency of toxin-inactivating deletions in the presence of intact antitoxin genes and CRP-S promoter (51%), coupled with the absence of any deletion that inactivates antitoxin but preserves toxin indicates that unopposed toxin is indeed harmful under some natural circumstances.

We have examined the *toxTA* operon from many angles and answered our initial question of why *toxA* knockout prevents competence in *H. influenzae,* but have raised new questions whose eventual answers we hope will give us greater insight not just into the *toxTA* system, but competence regulation in general.

## METHODS

### Bacterial strains, plasmids, and growth conditions

Bacterial strains used in this work are listed in Supp Table 1. *Escherichia coli* strain DH5 *α* [F80lacZ #(lacIZYA-argF) endA1] was used for all cloning steps; it was cultured in Luria-Bertani (LB) medium at 37°C and was made competent with rubidium chloride according to the method provided in the QIAexpressionist manual protocol 2 (Qiagen). When antibiotic selection was required, 100 µg/mL ampicillin and 50µg/mL spectinomycin were used.

*Haemophilus influenzae* cells were grown in sBHI medium (Brain Heart Infusion medium supplemented with 10mg/mL hemin and 2mg/mL NAD) at 37°C in a shaking water bath (liquid cultures) or incubator (plates). *H. influenzae* strain Rd KW20 (Alexander and Leidy 1951), the standard laboratory strain, was used as the wild type for all experiments. Mutant strains used in this study were marked deletion mutants in which the coding region of the gene was replaced by a spectinomycin resistance cassette, as well as unmarked deletion mutants derived from these strains; the generation of these mutant strains is described in Sinha et al. (2012). Specifically, we used an unmarked deletion of HI0659 (HI0659-), marked and unmarked deletions of HI0660 (HI0660::spec, HI0660-), and a marked deletion of the whole operon (HI0659/HI0660::spec). Knockout mutants of *crp* and *sxy* have been described previously (Chandler, 1992, Williams *et al.,* 1994)

*Actinobacillus pleuropneumoniae* cells were grown in BHI-N medium (Brain Heart Infusion medium supplemented with 100µg/mL NAD) at 37°C. *A. pleuropneumoniae* strain HS143 (Blackall *et al.* 2002) was used as the wild type for all experiments. Marked deletion mutants in which the gene of interest was replaced by a spectinomycin resistance cassette strains were generated for this study as described below. The HS143 genome region containing the homologs of the *Actinobacillus pleuropneumoniae serovar 5b strain L20* APL_1357 and APL_1358 genes, plus approximately 1 kb of flanking sequence on each side, was PCR-amplified, ligated into Promega pGEM-T Easy and transformed into *E. coli*. Plasmid regions containing APL_1357, APL_1358, or both genes were deleted from the pGEM-based plasmid by inverse PCR, and the amplified fragments were blunt-end ligated to the spectinomycin resistance cassette (Tracy *et al.,* 2008) from genomic DNA of a *H. influenzae comN*::*spec* strain (Sinha *et al.,* 2012). Plasmids linearized with ScaI were transformed into competent *A. pleuropneumoniae* HS143 and transformants were selected for spectinomycin resistance using 100µg/mL spectinomycin after 80 minutes of growth in nonselective medium.

### Generation of competent stocks

To induce competence, *H. influenzae* and *A. pleuropneumoniae* were cultured in sBHI or BHI-N respectively and transferred to competence-inducing medium MIV (Herriot et al. 1970) when they reached an optical density at 600nm (OD_600_) of approximately 0.25 (Poje and Redfield 2003). After incubation with gentle shaking at 37°C for a further 100 min (*H. influenzae*) or 150 min (*A. pleuropneumoniae*), cells were transformed or frozen in 16% glycerol at −80 °C for later use.

### Transformation assays

#### Transformation of MIV-competent cells

Transformation assays were carried out as described by Poje and Redfield (2003). MIV-competent *H. influenzae* or *A. pleuropneumoniae* cells were incubated at 37°C for 15 minutes with 1µg/ml DNA, then DNaseI (10µg/mL) was added and cultures were incubated for 5 minutes to ensure no DNA remained in the medium. *H. influenzae* cultures were transformed with MAP7 genomic DNA (Barcak *et al.* 1991), which carries resistance genes for multiple antibiotics, while *A. pleuropneumoniae* cultures were transformed with genomic DNA from an *A. pleuropneumoniae* strain with spontaneous nalidixic acid resistance (generated in this lab). Cultures were diluted and plated on both plain and antibiotic-containing plates (2.5ug/mL novobiocin for *H.influenzae* cultures, 20ug/mL nalidixic acid for *A. pleuropneumoniae* cultures) and transformation frequencies were calculated as the ratio of transformed (antibiotic-resistant) cells to total cells. For *A. pleuropneumoniae,* transformed cells were given 80 minutes of expression time in BHI-N before plating.

#### Time courses in rich medium

*H. influenzae* cells from frozen stocks of overnight cultures were diluted in fresh sBHI and incubated with shaking at 37°C. Periodically, the OD_600_ was measured, and at predetermined optical densities aliquots of the culture were removed and transformed with MAP7 DNA and plated as described above. **Bioscreen Growth Analysis:** The Bioscreen C apparatus (BioScreen Instruments Pvt. Ltd.) was used to measure growth. Cells frozen from overnight cultures were pre-grown at low density in sBHI, and 300µL aliquots of 100-fold dilutions were placed into 20 replicate wells of a 100-well Bioscreen plate. Wells at the edges of the plate were filled with medium alone as controls. Cells were grown in the Bioscreen at 37°C for 18 hours with gentle shaking, and OD_600_ readings were taken every 10 minutes. Readings were corrected by subtracting the OD_600_ measured for medium-only controls, and replicates for each strain were averaged at each time point to generate growth curves. Doubling times were calculated for each strain from the subset of time points that represents exponential growth phase, as determined by linearity on a semi-log plot of time versus OD_600_.

#### Competence growth and survival time course

Cells were grown in sBHI to a density of ∼2×10^8^ cfu/ml (OD600 = 0.075) and transferred to MIV. After 100 min (time for maximum competence development, an aliquot of each culture was diluted 1/10 into fresh sBHI for recovery and return to normal growth. A fraction of each culture was incubated in a shaking water bath, and aliquots of the initial and ‘recovery’ sBHI cultures were also grown and monitored in a Bioscreen incubator.

#### Cyclic AMP competence induction

*H. influenzae* cells in sBHI were incubated with shaking to an OD_600_ of approximately 0.05. Cultures were split and 1mM cAMP was added to one half. At an OD_600_ of approximately 0.3, aliquots were transformed with MAP7 DNA and plated as described above.

#### Phylogenetic Analysis

A nucleotide BLAST search (discontinuous MEGABLAST) and a protein BLAST search against translated nucleotide databases (tBLASTn) were used to identify homologs of the HI0659 and HI0660 genes (Altshul *et al*. 1990). Protein sequences found by the tBLASTn search were retained for analysis if they showed greater than 60% coverage and greater than 40% identity to the *H. influenzae* query sequence. For species with matching sequences in multiple strains, the sequence from only one strain was kept.

For species in which homologs of HI0659 and HI0660 were found next to one another, amino acid sequences of concatenated matrices were aligned by multiple-sequence alignment using MAFFT, version 7.220 (Katoh, 2013), run from modules within Mesquite version 3.02 (Maddison and Maddison 2015). The L-INS-I alignment method was used due to its superior accuracy for small numbers of sequences. After inspection of the alignments, poorly-aligning sequences were removed from the analysis, and alignment was repeated.

Phylogenetic trees were generated using the RAxML (Stamatakis 2014) maximum likelihood tree inference program, run via the Zephyr package of Mesquite. For each gene, 50 search replicates were conducted, using the PROTGAMMAAUTO option to allow RAxML to automatically select the best protein evolution model to fit the data. Since these trees were found to correspond exactly to a set of trees generated using the PROTGAMMAJTT model, this faster model was used to generate a majority-rules consensus tree from 1000 bootstrap replicates for each gene.

#### Analysis of natural deletions

181 publicly available *H. influenzae* genomes were downloaded from NCBI and the Sanger centre. (Supp table #???) Genomes were re-annotated using Prokka v1.11 (Seemann, 2014), and the pangenome was calculated using Roary v3.5.1 (Page *et al.*, 2015) with a minimum blastp threshold of 75. The *toxA* gene cluster in the pangenome was identified by finding the gene cluster that contained the *toxA* gene from Rd KW20, and the *hicA* cluster was identified by finding the gene cluster that contained the *hicA* gene from PittAA. 2300 bp genome sequences centered on *toxA* and/or *hicA* were extracted from all *H. influenzae* genomes containing recognizable *toxA* and/or *hicB* genes, and aligned by the MAFFT server. For strains that lacked recognizable *toxA* or *hicB*, sequences adjacent to the genes that normally flanked each operon were extracted. Ka/K_s_ and pairwise distance were calculated for each gene using SeqinR v 3.4-5 (Charif and Lobry, 2007) with codon aware gene alignments were made using Prank (v.100802).

### RNA-seq analysis

#### Sample Preparation

Cell cultures of *H. influenzae* strain Rd, Δ*crp* and Δ*sxy* derivatives, and Δ*toxTA* mutants were grown in sBHI to an OD_600_ of 0.2 – 0.25, then transferred to MIV. Aliquots of cells were removed just prior to transfer to MIV, and after 10, 30, and 100 minutes in MIV, and immediately mixed with Qiagen RNAprotect (#76526) to stabilize RNA. Cells were pelleted and frozen, and RNA was later extracted from thawed pellets using the Qiagen RNeasy Min-elute Cleanup Kit (#74204). Contaminating DNA was removed with Ambion Turbo DNase (#AM2238), and ribosomal RNA was depleted using the Illumina Ribo-Zero rRNA Removal kit (#MRZMB126). Sequencing libraries were prepared using TruSeq mRNA v2 library preparation kit, according to manufacturer’s instructions (Illumina).Libraries were pooled and sequenced on a HiSeq 2500, generating paired-end 100 bp reads.

#### Data Analysis Pipeline

FASTQ files were analysed using the FASTQC tool (Andrews, 2015) to confirm read quality. Reads were aligned to the *H. influenzae* Rd KW20 reference genome sequence using the Burrows-Wheeler Alignment tool (BWA) algorithm bwa mem (Li and Durbin, 2009). Differential expression analysis was performed using the DESeq2 package, v.1.6.3 (Love *et al.,* 2013). Specifically, the function DESeqDataSetFromMatrix()was used to generate a dataset to compare reads from each mutant strain reads from the wild-type control based on their strain, sample time point, and the interaction between the two parameters. The function DESeq() was called to determine which genes were differentially expressed based on these parameters, using p-values adjusted for a B-H false-discovery rate (Benjamini and Hochberg, 1995) of 0.1 as a cut-off to determine significance, after normalizing total read counts and variances.

## ACKNOWLEDGEMENTS

We thank Lauri Lintott for helpful discussions, Charles Thompson for the use of the BioScreen, and Anni Zhang and Yvonne Yiu for technical assistance. This work was supported by funding from Canadian Institutes of Health Research to RJR, and an NIH F32 AI084427 grant to JCM. Sequencing work was performed at the Sequencing and Bioinformatics Consortium at the University of British Columbia, supported by grants from NASA and the faculty of Pharmaceutical Sciences.

